# A Self-attention Graph Convolutional Network for Precision Multi-tumour Early Diagnostics with DNA Methylation Data

**DOI:** 10.1101/2022.02.11.480173

**Authors:** Xue Jiang, Zhiqi Li, Aamir Mehmood, Heng Wang, Qiankun Wang, Yanyi Chu, Xueying Mao, Jing Zhao, Mingming Jiang, Bowen Zhao, Guanning Lin, Edwin Wang, Dongqing Wei

**Affiliations:** School of Life Sciences and Biotechnology, Shanghai Jiao Tong University, Shanghai 200030, China; Department of Biochemistry and Molecular Biology, Medical Genetics, and Oncology, Cumming School of Medicine, University of Calgary, Calgary, Alberta, Canada; International School of Cosmetics, School of Perfume and Aroma Technology, Shanghai Institute of Technology; School of Biomedical Engineering, Shanghai Jiao Tong University, Shanghai 200030, China

**Keywords:** Self-attention mechanism, Graph convolutional network, Key methylation sites, Multi-tumour early diagnostics

## Abstract

DNA methylation data-based precision tumor early diagnostics is emerging as state of the art technology, which could capture the signals of cancer occurrence 3∼5 years in advance and clinically more homogenous groups. At present, the sensitivity of early detection for many tumors is about 30%, which needs to be significantly improved. Nevertheless, based on the genome wide DNA methylation information, one could comprehensively characterize the entire molecular genetic landscape of the tumors and subtle differences among various tumors. With the accumulation of DNA methylation data, we need to develop high-performance methods that can model and consider more unbiased information. According to the above analysis, we have designed a self-attention graph convolutional network to automatically learn key methylation sites in a data-driven way for precision multi-tumor early diagnostics. Based on the selected methylation sites, we further trained a multi-class classification support vector machine. Large amount experiments have been conducted to investigate the performance of the computational pipeline. Experimental results demonstrated the effectiveness of the selected key methylation sites which are highly relevant for blood diagnosis.

## 1. Introduction

The world health organization (WHO) of the internal agency for research on cancer (IARC) states that there were 19.29 million cancer cases have appeared and 9.96 million cancer deaths in 2020 [1]. The world’s cancer situation is grim. With the aging and rapid growth of the population, cancer incidence and the number of deaths continue to rise. While much of current research is focused on developing new therapeutics, studies have shown that early detection has the potential to reduce both treatment cost and mortality rates from cancer by a significant amount.

Early cancer diagnosis offers the opportunity to identify tumors when cures are more achievable, and treatment can be less morbid. However, most cancer types currently lack an effective non-invasive early screening option [2]. Effective screening paradigms exist only for a small subset of cancers and focus on single cancer type. In recent years, DNA methylation (5-methylcytosine, 5mC) in cell-free DNA (cfDNA) has been successfully detected in clinical samples by a range of genome-wide approaches, demonstrating high clinical potential in cancer early diagnosis, prognosis, and/or treatment response.

DNA methylation is a primary epigenetic mechanism involving covalent modification of cytosine bases by adding a methyl group at the 5’ position. DNA methylation plays a critical role in the normal development and regulation of many cellular processes [3], with DNA methylation profiles being cell type and tissue-specific [4]. DNA methylation occurs before the tumor and is an essential mechanism of it. A global loss of DNA methylation (hypomethylation) accompanied by increased de novo DNA methylation (hypermethylation) of CpG-rich regions can be implicated in many tumors [5, 6]. Generally, the DNA of dead tumour cells can be released and circulates in the blood stream, termed as circulating tumor DNA (ctDNA). The ctDNA can be detected in the blood to enable early tumor diagnosis of 3∼5 years (ultra-early) before its formation. DNA methylation based tumor diagnostics is currently emerging as state of the art technology [7-10].

Precision early tumor diagnostics is crucial for patients. Studies have shown that early tumor detection can significantly increase a patient’s 10-year survival rate (from 30% to 90%). The 5-year survival rate for patients with early-stage can be up to 91% if treated with the early intervention[11-13]. DNA methylation is particularly suitable for personalized early tumor diagnostics for three reasons: 1) a large number of researchers have identified characteristics differences in the DNA methylation profiles between subgroups of patients [14-18], demonstrating the power of DNA methylation for analyzing epigenetic heterogeneity, 2) since the cancer methylome is a combination of both somatically acquired DNA methylation changes and characteristics reflecting the cell of origin, it is especially suitable for molecular classification of tumors [19-22], 3) it has been convincingly shown that DNA methylation profiling is highly robust and reproducible even from small samples and poor quality materials [23]. Although 5mC biomarkers based on plasma cfDNA are sensitive for early tumor detection, in large-scale population screening, such a multi-tumor detection approach would require high specificity and useful clinical sensitivity to limit the scope, cost, and complexity of evaluating asymptomatic patients.

Nowadays, the sensitivity of the early (stage I) diagnostics of many tumors, including breast, esophageal, kidney, lung, prostate, and uterine cancers, is about 30%, which needs to be significantly improved. Existing studies of tumor diagnostics with DNA methylation mainly include two steps. Firstly, information regions, including differentially methylated CpG sites and cancer-related genomic regions reported in the literature, are determined. Then targeted panel is made for in-depth sequencing. Secondly, regression models often used to fit the methylation beta value data of the selected sites. The bottlenecks in the current research can be shown in three aspects. First, as we all know that cancer is a genome-wide disease, key methylation sites determined with the knowledge-driven way would restrict the understanding of disease pathological mechanisms. Second, the linear classification accuracy depends heavily on the selected methylation sites, limiting its generalization and extensibility. Third, the existing bioinformatic tools have not fully utilized the whole-genome methylation information to conduct early tumor diagnostics.

As the development of sequencing and chip technology, it provides great opportunities and challenges for precision early diagnostics of many tumors. Therefore, a methodology for DNA methylation analysis, which could effectively use whole-genome information and improve the early diagnosis sensitivity, is urgently needed. Based on the above analysis, advanced graph neural network model in the field of artificial intelligence is suitable for modeling the high dimensional data, improving the robustness and accuracy of early tumour detection. In this study, a computational framework of a self-attention graph convolutional network (SAGCN) model and a multi-class classification support vector machine was designed to distinguish the 11 most common cancers with the DNA methylation data. On the one hand, the self-attention graph convolutional network learned methylation sites interaction networks in an end-to-end way, automatically obtained key methylation sites (namely, differentially expressed methylation sites, DEMs); on the other hand, the computational framework improved the stability and accuracy of prediction while the sequencing depth is insufficient. Collectively, the framework proposed here advances multi-tumor diagnostics with robust and higher accuracy of early tumor diagnostic.

## 2. Materials and Methods

### 2.1 Study design and data preparation

We propose a workflow based on SAGCN to conduct feature selection and sample classification with the DNA methylation data. This workflow has four parts. Part 1 is data preparation, preprocessing, and basic filtering. Part 2 involves the training process with SAGCN. The clinical sample classification with SVMs is done in part 3. And finally, Part 4 is performance evaluation. All the steps are diagrammatically represented in Figure 1.

**Fig. 1.**
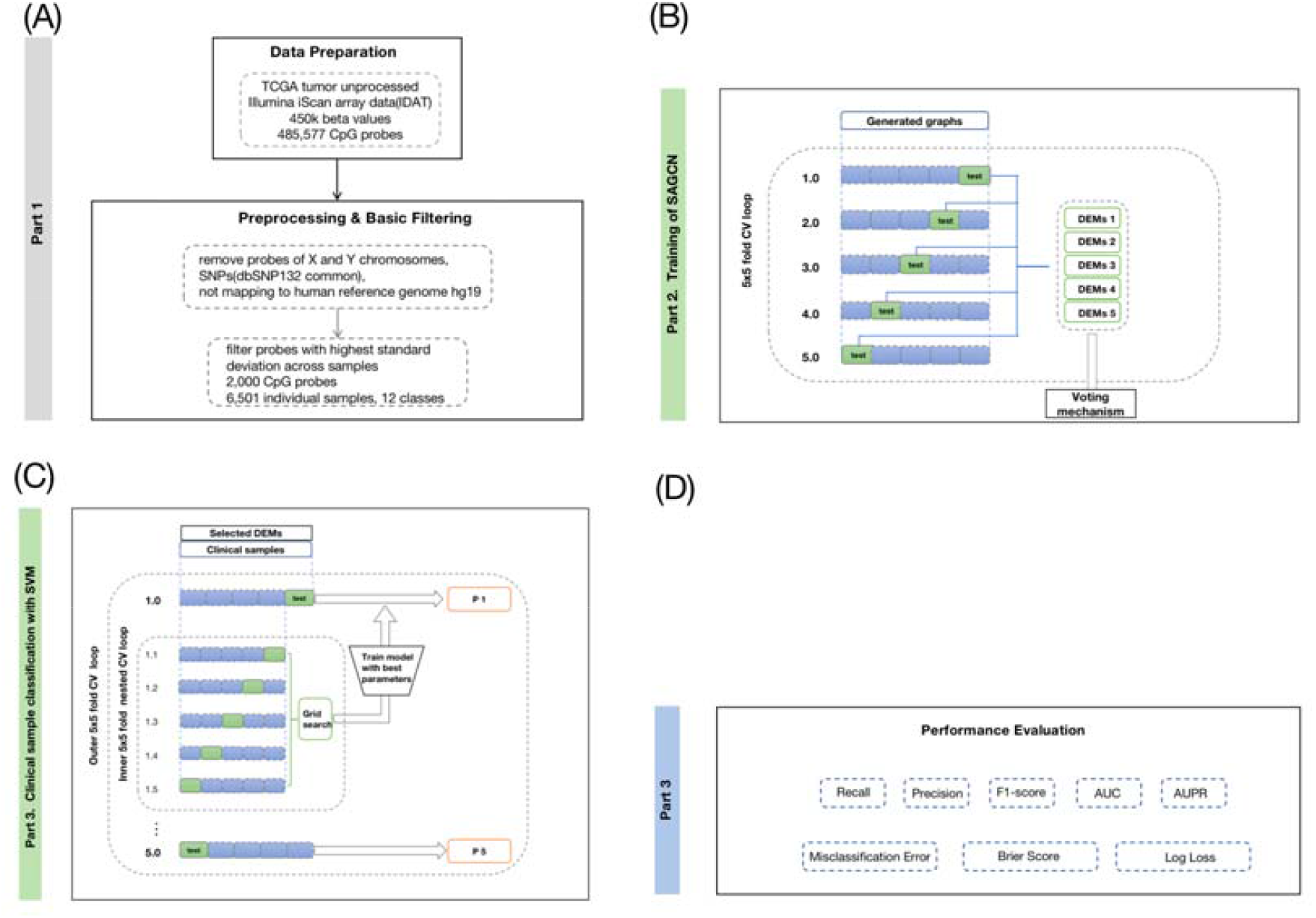
The pipeline of the self-attention graph convolutional network based computational framework.

From The Cancer Genome Atlas National Cancer Institute Genomic Data Commons (TCGA NCI GDC) Repository (https://portal.gdc.cancer.gov/repository), we collected 450k DNA methylation data from 6557 samples, including 11 types of primary tumor site, such as bladder, brain, breast, bronchus and lung, cervix uteri, corpus uteri, kidney, liver and intrahepatic bile ducts, prostate gland, stomach, and thyroid gland. The samples include 5905 primary tumor sites, 596 solid tissue normal samples, 37 recurrent tumor samples, 17 metastatic samples, 2 new primary samples. Since the sizes of recurrent tumor samples, metastatic samples, and new primary samples are too small, we only utilize primary tumor samples and solid tissue normal samples for further analysis. This study divides the samples into 12 classes, all solid tissue normal samples and 11 types of tumor samples. The distribution of these samples is shown in Supplementary Figure S1.

The detailed patient demographics and clinical characteristics can be found in Supplementary Table 1. For details on how to prepare the 450k DNA methylation tumor samples from TCGA, please see the GitHub repository (https://github.com/lizhiqi0506/GNN_CancerPreDiagnosisWithMeth), and to download the source data, visit the NCI GDC Legacy Archive (https://gdc-portal.nci.njh.gov/legacy-archive).

Illumina Human Methylation BeadChip arrays are a popular tool to measure genome-wide single-nucleotide CpG site methylation levels [17]. The DNA methylation data is based on genome-wide quantitative measurements of DNA methylation at 485,577 CpG sites using Illumina Human Methylation 450 BeadChip technologies (450k; Illumina). The 450k BeadChip provides >98% coverage of reference sequence genes and 96% of CpG islands. The beta value represents the average methylation fraction (AMF), computed for each genomic region by summing the number of observed cytosines at all covered CpG sites and dividing by the total sequencing depth at all covered CpG sites in each region. For each methylation locus, the amount of methylated DNA is denoted as Meth, and the amount of unmethylated DNA is denoted as Unmeth. The beta value is denoted as Meth/(Unmeth + Meth). The beta value is further used in downstream analysis.

The matrix 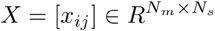 is used to represent the DNA methylation beta value, in which *N*_*m*_ is the number of methylation sites, *N* _*s*_ is the number of samples. To reduce the complexity of downstream analysis and improve the performance of the computational method, preprocessing and basic filtering steps are needed.

Step 1. filter out samples with uncompleted site covers.

Step2. remove probes of X and Y chromosomes, probes containing single-nucleotide polymorphisms (dbSNP132Common), and probes not mapping uniquely to human reference genome 19. Finally, 396,013 methylation sites were left.

Step 3. 2000 most variably methylation sites were further selected according to the highest standard deviation across all samples.

Step 4. then, a principal component analysis (PCA) was conducted. The first 100 principal components were then used as input data for the t-Distributed Stochastic Neighbour Embedding (t-SNE, Rtsne package version 0.11, https://github.com/mwsill/mnp_training/blob/master/tsne.R) for clustering analysis [26, 27]. The cluster result is shown in Fig. 2. From Fig. 2, we can know that the samples can be distinguished clearly. However, we need a high performance machine learning method to automatically select key methylation sites that can classify samples more accurate and stable.

**Fig. 2.**
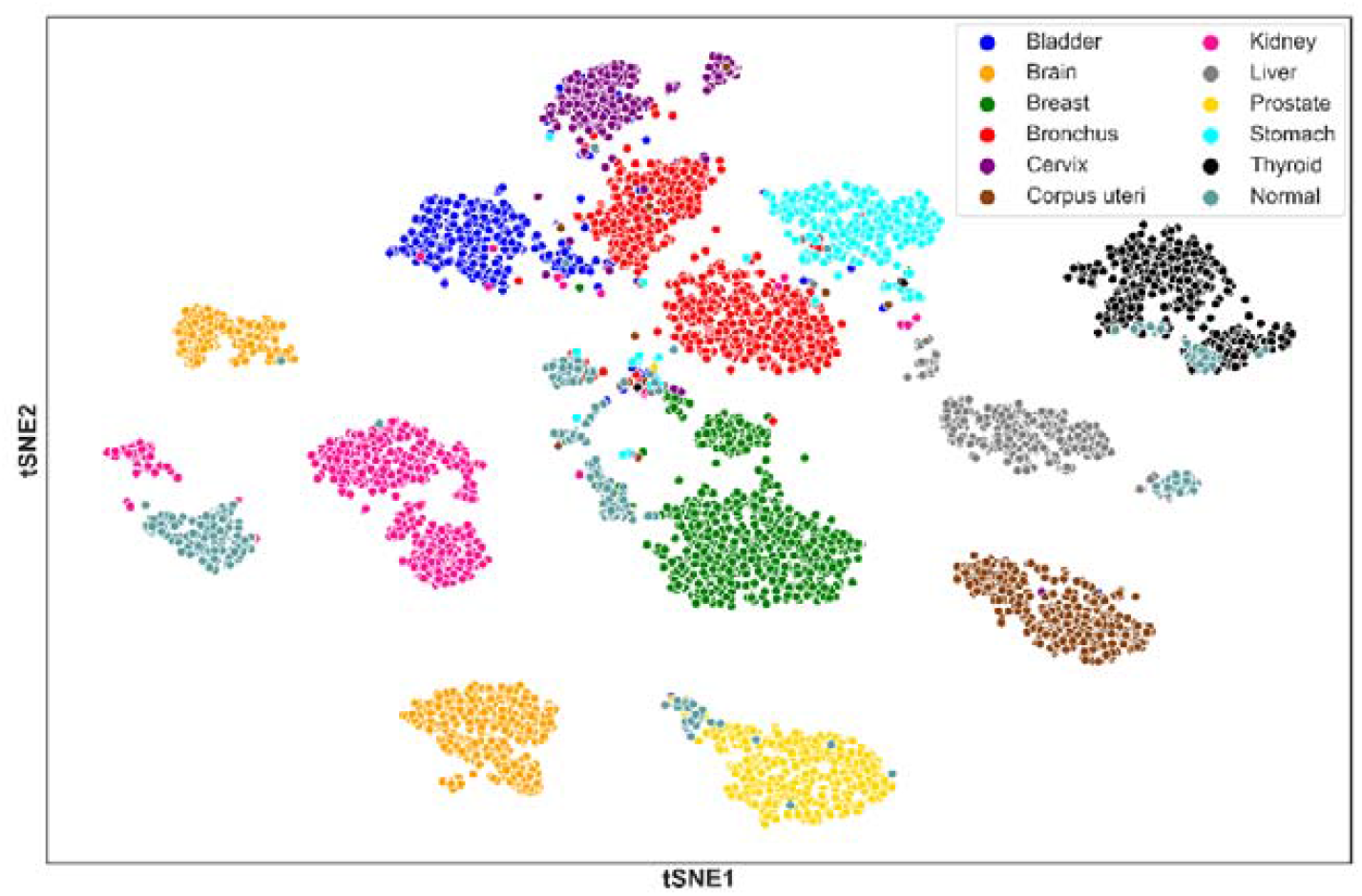
t-SNE visualization of 6501 samples analyzed by PCA, colored by clusters, and labeled by the sample type.

### 2.2 Self-attention graph convolutional network

#### 2.2.1 Construction of methylation sites interaction graph

This study used weighted gene co-expression network analysis (WGCNA) to construct a methylation site interaction graph. Let *G(V,A,X)* represents a methylation site interaction graph of one class, where represents the methylation sites in the network, 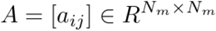 is the adjacency matrix, 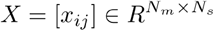 is the original feature matrix, *N*_*m*_ is the number of methylation sites, *N*_*s*_ is the number of samples. It should be noted that, in the network construction, the interaction relationships between methylation sites were determined by the following criterion:

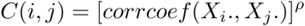

where *corrcoef(*·, ·) is a function that calculates the Pearson correlation coefficient between edges. The power for bladder, brain, breast, bronchus, cervix, corpus, kidney, liver, prostate, stomach, thyroid, and normal is 5, 3, 8, 6, 3, 4, 7, 5, 2, 5, 3 and 10 respectively. So most weights of the edges are small values (Supplementary Figure S2 shows the weight distribution for each network). Therefore, we set a threshold 0.01 to determine whether there is an interaction between two methylation sites. Further more, we conducted significance analysis of the relationships in each network. The P-values for the weights are far less than 0.05, indicating a significant correlation of the relationships in the network.

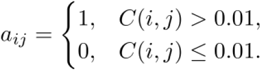

It should be noticed that, we used a subsampling strategy to deal with the small sample size and label imbalance problem, the detailed clinical information statistic can be seen in Supplementary Table 1. We randomly sampled 200 samples from the corresponding dataset with replacement for each class. Through the above strategy, 80 graphs were created for each class (totally 80 x 12 = 960 graphs).

#### 2.2.2 Graph convolutional network

Deep learning has made great progress in computer vision and natural language processing. Meanwhile, in the real world, most issues cannot be described by Euclid field data but can be abstracted as graphs easily. To perform deep learning based on graph data, different kinds of graph neural networks (GNNs) were presented. Graph convolutional network (GCN) is one of the most widely used GNNs. Graph convolutional operation can be defined in either the spectral or non-spectral domain. Spectral approaches perform convolution on a graph by redefining the convolution operation in the Fourier domain. The classical spectral graph convolution method is formulated as below:

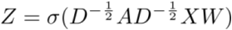

Where 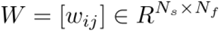 represents weight matrix, *N*_*f*_ is the updated dimensionality of output features, represents adjacency matrix for methylation sites interaction graph, *D* is the diagonal matrix of *A* with 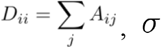 is a nonlinear activation function, 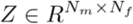 is feature matrix after convolution.

#### 2.2.3 Self-attention graph pooling

Attention mechanisms help to focus on those essential features and lose sight of trivial features, which is very useful in classification and data mining. This study proposed a graph pooling method with an attention mechanism (SAGPooling). The attention mechanism is realized by graph convolution. For a methylation sites interaction graph with *N*_*m*_ nodes, the self-attention score 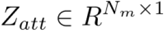 is defined and calculated as

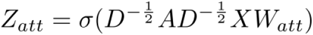

Where 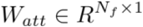 is the only parameter of SAGPooling layer. Both node properties and graph topology are considered by using graph convolution to obtain self-attention scores. The hyperparameter of pooling ratio is used to determine the nodes selected during the step. We ranked the nodes in descending order based on the value of *Z*_*att*_ and selected top *kN*_*m*_ nodes.

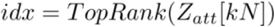

Where *Top Rank* is a function that returns the indices of the top [*kN*] values. For an input graph *G(V,A,X)*, the output graph *G*′(*V*′,*A*′,*X*′) can be obtained as:

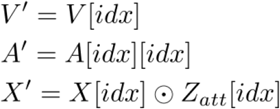

Where [*idx*] is an indexing operation, ⊙ is the broadcasted elementwise product.

#### 2.2.4 Self-attention graph convolutional network model

We proposed a graph neural network mixed with GCN layers and SAGPooling layers (SAGCN). The structure of SAGCN can be divided into two parts. The first part is used for node selection and graph embedding. It contains 3 sequential modules, with a GCN layer followed by a SAGPooling layer in each module.

The activation function of the GCN layer is ReLU. For an input graph *G*^(0)^ (*V*^(0)^, *A*^(0),^ *X*^(0)^), in the first module, we get *G*^(1)^ (*V*^(0)^, *A*^(0),^ *X*^(1)^) after the GCN layer. And then, SAGPooling is performed on *G*^(1)^ :

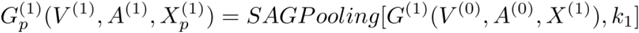

Where *SAGPooling*[.,.] operation of SAGPooling layer and *k*_1_ is the pooling rate of the first module’s SAGPooling layer. And then 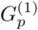 will be the input of the second module. Finally, after all the three modules processing, we get 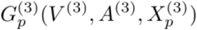 with *N*_*d*_= *N*_*m*_*k*_1_*k*_2_*k*_3_ methylation sites left, *k*_2_ and *k*_3_ are the pooling rate of SAGPooling layers of the second and third module, respectively. Finally, *V*^(3)^ is the differentially expressed methylation sites (DEMs) selected from the input graph by the algorithm.

Next, we perform scatter-max and scatter-mean on 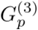 to get a graph embedding (GE). Scatter-max operation is to pick the maximum element in each row as the feature of the corresponding node. Scatter-mean operation calculates the average value of each row as the feature of the corresponding node. GE is generated by concatenating the two vectors and then serves as the input of the second part.

The second part of the model mainly functions as a classifier, consisting of a CNN layer and a multilayer perceptron (MLP) with one hidden layer. Finally, for the computational convenience of the loss function, a log softmax is computed on the output of MLP as the prediction score. The SAGCN model is illustrated in Figure 3.

**Fig. 3.**
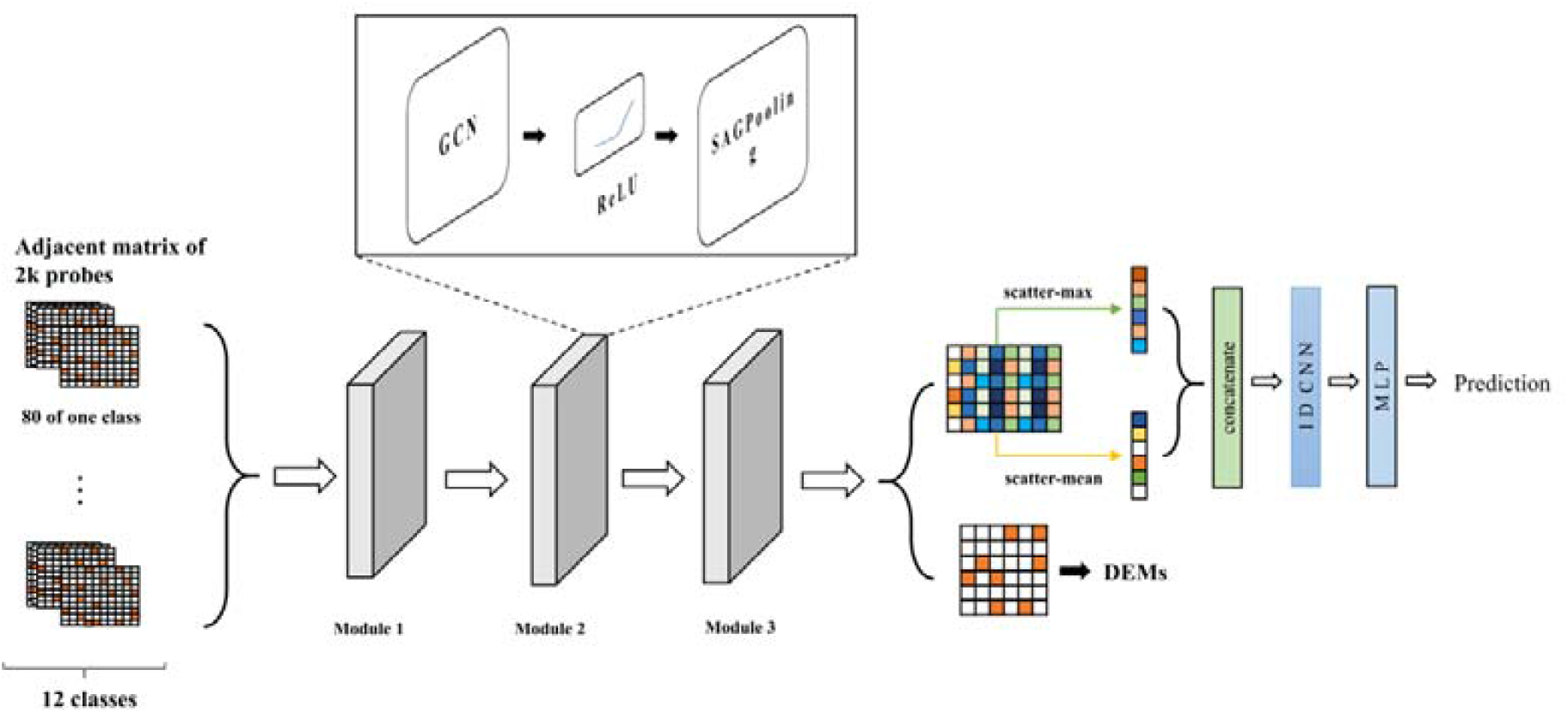
The workflow of the proposed SAGCN model.

### 2.3 Selecting DEMs

To obtain a tissue-specific methylation pattern, we first construct interaction relationships between these methylation sites for each class. Then, those graphs are used to train the SAGCN, in which graph convolutional layer was used to aggregate features, and a self-attention mechanism (SAGPooling) was used to extract the DEMs.

For the input graph, we get the methylation sites *V*^(3)^ from the first part of the SAGCN model. For different input graphs, theoretically, there is a minor change in the selected *V*^(3)^. Therefore, to get a convergent and robust DEMs set, we utilized a voting mechanism to determine the final selection that shows up most frequently for all input graphs.

### 2.4 Clinical sample classification with SVM

To verify the validity of the selected DEMs, we further trained a multi-class classification support vector machine model with a nested 5x5 cross validation loop. The trained SVM model can be used to conduct clinical sample classification.

### 2.5 Performance evaluation

In this study, eight criteria were used to evaluate the performance of the model, including recall, precision, F1-score, area under the receiver operating characteristic curve (AUROC), an area under the precision-recall curve (AUPR), misclassification error (ME), Brier score (BS), and log loss (LL).

The formulas to compute the recall, precision, F1-score, and ME are as below:

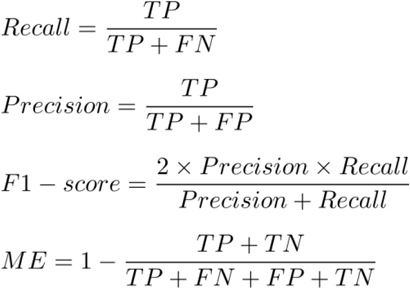

Where TP represents true positive rate, TN represents true negative rate, FP represents false-positive rate, and FN represents false negative rate in the predicting results. Then, the macro averaged recall, macro averaged precision, macro averaged F1-score, and macro averaged ME to evaluate the overall model performance of multi-classification. Moreover, macro averaged values of AUC and AUPR are also calculated. It should be noted that the averaged macro evaluation criterion is the average of the evaluation criterion measured in each of the 5x5-fold CV (outer) test sets.

BS is a proper scoring rule that measures the accuracy of probabilistic predictions of mutually exclusive classes. It is applicable to multiclass prediction and is defined as the quadratic difference between the assigned probability and the value (1, 0) for the class:

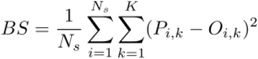

where *N*_*s*_ is the number of samples and *K* is the number of classes. *P*_*i,k*_ is calibrated estimation probability that observation ***i*** belongs to class ***k*** and *O*_*i,k*_ is the actual class of sample ***i***.

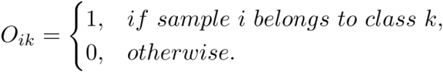

Besides, LL is extensively used to assess probability estimates of predictor models. BS encourages predicted probabilities and true labels to lie close to each other, whereas LL does not. However, extensive empirical testing stressed LL’s favorable local property that it will always assign a higher score to a higher probability estimate for the correct class. In contrast, BS can perform poorly in this regard. Thus we also used LL to evaluate model performance. A multiclass extension of log loss is shown below:

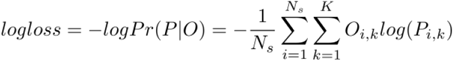

## 3. Results

### 3.1 Performance of SAGCN

Large amounts of experiments have been performed to investigate the performance of the computational framework. DEMs could be extracted through the pooling rate in part one of SAGCN, making sure robust and good classification performance. To evaluate classification performance, 5x5-fold cross-validation was implemented. And macro average values of recall, precision, F1-score, AUC, AUPR, ME, BS, and LL are calculated. We hold the pooling rate of the first and the second SAGPooling layers, i.e., *k*_1_ = *k*_2_ = 0.5, and changed the value of *k*_3_ to determine the number of finally selected DEMs, denoted as. The classification performance with different parameters *k*_3_ is shown in Table 1. We can observe that the performance of SAGCN becomes better then worse with the increase of *k*_3_, and it achieves the best performance when *k*_3_ = 0.5. The recall, precision, F1-score, AUC and AUPR are 0.9917, 0.9918, 0.9917, 0.9998, and 0.9995, respectively. The AUC and AUPR are greater than 0.99, indicating that the proposed method has an outstanding classification ability. The ROC and PR curves with *k*_3_ = 0.3 are shown in Supplementary Figure S3, demonstrating the outstanding performance of the SAGCN.

There are minor changes to the selected *V*^(3)^ for each cross validation fold during the training process. Therefore, to get a convergent and robust DEMs set, we used a voting mechanism to determine the final selection that frequently shows up. We conduct experiments for sample classification with different *k*_3_ values to assess the selected DEMs. The experimental results are shown in Table 2, according to which we selected 350 DEMs for downstream analysis. The extracted DEMs and related annotations can be seen in Supplementary Table 2. To illustrate the effectiveness of selected DEMs, we draw the cluster heatmap (seaborn, python package, version 0.10.1) with their beta values (Supplementary Figure S4), where the rows represent the DEMs and the columns represent samples that are sorted according to the class order. It illustrated that different blocks are formed among different classes, indicating the effectiveness of selected DEMs.

Furthermore, to test the classification performance of SAGCN for each class, the mean values of the basic metrics for each class on 5 folds are shown in Table 3. For the 11 kinds of tumor samples, the evaluation metrics are all greater than 0.95, except the precision of the cervix with a value of 0.93, which suggests that nearly 6.73% of the cervix classified as tumor samples are misclassified. Additionally, normal samples’ recall, Precision, F1-score, AUC, and AUPR are relatively small. The Precision is only 0.8925, which means that nearly 10.75% of normal samples are misclassified as tumor samples. The reason is that, firstly, the normal samples are from different tissues, so they are characterized by tissue-specific DNA methylation features, which may have an intersection with tissue-cancer-specific DNA methylation sites, leading to a high misclassification. Secondly, from supplementary Figure S4, we can see that the expression of selected DEMs is closer between Thyroid gland tumor samples and normal samples, which easily causes a misclassification between thyroid gland tumor samples and normal samples.

To verify the effectiveness of DEMs selected by SAGCN, we further trained a multi-class classification SVM, namely SAGCN+SVM. More importantly, we checked and verified the classification performance of SAGCN + SVM for tumors of stage I and stage II with the selected DEMs. The results are shown in Table 4 and Table 5, respectively. There are 7 kinds of tumor with stage I and stage II in the dataset, totally 2615 samples. The recalls, i.e., sensitivity, of them are all larger than 0.95, illustrating the effectiveness of the computational pipeline and the selected DEMs. In brief, the SAGCN computational framework can effectively prioritize DEMs, which could get super classification performance on nearly all kinds of tumour samples.

### 3.2 Comparisons with classic machine learning models

We compared it with some classic machine learning algorithms based on the original 2000 methylation sites. This included random forest classifier (RF), extremely randomized trees (ERT), decision tree (DT), Gaussian naive Bayes (GNB), and SVM. Table 6 tabulates the prediction results where we can see that the performance of SAGCN + SVM is better, indicating the effectiveness of SAGCN and the selected DEMs.

For the reason of multi-classification, the classification of each class is a binary classification on an unbalanced dataset. In this term, the high accuracy of each class is less believable. Inversely, Recall, Precision, F1-score, AUC, and AUPR can genuinely reflect classification performance.

### 3.3 Comparisons with the state-of-the-art methods

To further prove the superiority of SAGCN+SVM, we compared it with two state-of-the-art methods proposed after 2020. One is a graph convolutional network-based method for predicting classification (CGATCPred); another is a hybrid graph convolutional network for predicting cancer drug response (DeepCDR). The following experiments are carried out under the same experimental conditions, including 5-fold cross-validation, random seed, and data partitioning strategy. The three methods can all achieve good performance (Table 7). The statistical test between SAGCN + SVM and the other two methods is 0.08 and 0.27 for AUC, respectively, indicating the sample classification performance of the proposed model is compared able to the state-of-the-art methods.

### 3.5 External validation with tissue DNA methylation data

To test the stability and extendibility, we applied this computational framework to external data from NCBI Gene Expression Omnibus (GEO) database, under accession GSE90496 [17], GSE155207 [30], GSE158075 [31], and GSE164988. The brain tumors DNA methylation dataset (GSE90496) includes 2801 samples from which 294 (294/2801) were selected, the disease stage of which is initially operated, the age distribution in the range of 25 to 70, and the WHO grade is I, II or III. A fumarate hydratase-deficient renal cell carcinoma dataset (GSE155207) was used as kidney tumour validation. The EPIC array methylation data of 20 primary tumor tissues was selected to conduct a validation experiment. Besides, the Human Methylation450-based DNA methylation analysis of paired samples of bronchoscopic biopsy specimens either from the tumor side or the contralateral tumor-free bronchus in 37 patients with a definite lung cancer diagnosis (GSE158075) was used as a validation dataset for bronchus and lung cancer. Moreover, 10 samples with histopathology diagnosed as adenocarcinoma from GSE164988 were used for stomach cancer validation.

Performance evaluation of SAGCN + SVM on external validation datasets is shown in Table 8. The classification performance for brain and kidney tumors can achieve 100% precision. Moreover, the classification accuracy for bronchus & lung, and stomach tumors can achieve more than 94%. It can also achieve 80% for normal samples. The results indicate that the proposed computational framework is robust, generalizes well, and can be easily applied to other external methylation datasets to get a well-performing classification.

### 3.6 External validation with cell-free DNA methylation data

Recent analyses of circulating cell-free DNA suggest that approaches using tumor-specific alterations may provide new opportunities for early diagnosis. But detecting such alterations may be challenging because the total number of alterations may be scarce in individuals with low tumor burden, hence not all patients have noticeable changes [33]. This study has developed an end-to-end deep learning framework to realize ultra-sensitive in common cancers. The genome-wide methylation approach has leveraged the discovery of inherited risk loci for cancer detection, the inherited risk loci are often involved in tissue-specific DNA methylation.

Methylation features of cfDNA, such as CpG sites, contain information about recent events in the body, which can be used to detect and localize cancer [34-36]. cfDNA molecules from loci carrying tissue-specific methylation patterns can be used to identify cell death in a specific tissue. Consequently, methods based on identifying CpG site markers or exploring the methylome could be used to accurately recognize cfDNA released by cells and tissues in various human cancer. However, cfDNA is not uniformly distributed across the genome but rather is differentially sheared and located depending on chromatin organization, gene expression, tissue of origin, epigenetic marks, and mechanism of release among other factors [37-40]. Identifying the cell and tissue of origin of the cfDNA fragment could lead to localization of cancer of unknown primary with liquid biopsy.

Therefore, we further used cell-free DNA methylation data from the GEO database under accession number GSE157272 to conduct external validation. First, blood is collected from the patient individuals. cfDNA is extracted from plasma, processed into sequencing libraries, examined by WGS, mapped to the genome, and analyzed to determine cfDNA fragmentation profiles across the genome. 22 samples from GSE157272 with disease stages as high-grade prostatic intraepithelial neoplasia tissue, indolent prostate cancer tissue, and aggressive prostate cancer tissue were used to conduct prostate cancer validation.

In this study, we first extracted the 350 DEMs with SAGCN. Then multi-class classification was conducted with SVM. The extracted DEMs and related annotations can be seen in Supplementary Table 2. From Supplementary Table 2, we can know that there are 85.2% methylation sites among all annotated sites located in protein coding region, indicating that can be easily detected in cell free DNA methylation data. Experimental result with GSE157272 is shown in Table 9. The result indicates that the proposed computational framework is robust, generalizes well, and can be easily applied to conduct a blood-based non-invasive liquid biopsy.

## 4. Conclusion and Discussion

Cancer diagnosis is complex because most malignant tumors present with long periods of latency and lack of clinical presentation at early stages. Nevertheless, epigenetics provides a molecular link between genetic programming and environmental signals. Such changes in the DNA methylation and the somatic genomic DNA from the tumor tissue of origin are highly consistent in many disease models. Thus, DNA methylation plays an important role in the process of DNA expression and precision tumor early diagnostics, emerging as state-of-the-art for molecular tumor recognition.

However, at present, the sensitivity of primary diagnosis for multiple cancers, such as tumour of breast, lung, prostate, uterine, esophageal and kidney is about 30%, and the diagnostic technology needs to be improved. Besides, for cancer-specific methylation sites identification based on DNA methylation data, the existing computational tools cannot fully utilize the genome-wide methylation information, and the computational tools have much room for improvement. Finally, it is costly and time-consuming to make a ultra-deep sequencing for the target panel, which is decided based on preliminarily screened cancer-specific methylation sites.

To address these challenges and issues, we present a genome-scale computational framework for precision multi-tumour early diagnostics. First, methylation interaction graphs are properly constructed. Then, a self-attention graph convolutional neural network model is proposed to extract the DEMs based on those graphs and conduct sample classification based on the extracted methylation sites. To improve the sample classification accuracy and robustness with the selected DEMs, we further designed SVM with inner 5-fold nested cross-validation for input sample classification. Large amounts of experiments have been conducted and have shown the effectiveness and robustness of the framework performance compared with either classic machine learning methods or state-of-the-art methods. External validation with tissue DNA methylation data and cell-free DNA methylation data also illustrated the effectiveness of the selected DEMs, indicating that the proposed computational framework is easily applied to other external methylation datasets to get well-performing tumor classification and tissue origination. In should be noticed that, in the external validation, the accuracy of several tumor types is large than 91%, which means that both the DEMs and computational framework can be applied to clinical practice.

Nevertheless, there also two limitations of this study. First, the DNA methylation data used in this research is 450k, while there 30 million CpG sites in the whole genome. Second, the research is retrospective study, while prospective study is more helpful for the detection sensitivity improvement of natural population. In the future study, we plan to conduct prospective methylation sequencing research to develop biomarkers of multi-omics and multi-sites, improving the stability and specificity of early tumor diagnostics.

## Acknowledgements

The study is supported by grants from the National Science Foundation of China (Grant No. 32070662, 61832019, 32030063), the Science and Technology Commission of Shanghai Municipality (Grant No.: 19430750600), as well as SJTU JiRLMDS Joint Research Fund and Joint Research Funds for Medical and Engineering and Scientific Research at Shanghai Jiao Tong University (YG2021ZD02). The computations were partially performed at the Pengcheng Lab. and the Center for High-Performance Computing, Shanghai Jiao Tong University.

## Key Points

- We have designed a SAGCN to learn methylation interaction networks in an end-to-end way, which automatically extracted DEMs.
- Through the pipeline of SAGCN for feature selection and SVM for classification, we achieved precision multi-tumor early diagnostics in silicon-based simulations.
- The computational pipeline would significantly contribute to tumor classification and is highly relevant for blood diagnosis.

## Data available

The complete methylation data required for the construction and training of the classifier have been deposited in TCGA. The external validation data can be downloaded from the NCBI Gene Expression Omnibus (GEO), under accession number GSE90496, GSE155207, GSE158075, GSE164988, and GSE157272.

## Code available

All the codes used in this research are complied with Python programming language. The codes, some important data files and more detailed description are available at https://github.com/lizhiqi49/SAGCN.

**supplementary Figure S1.**
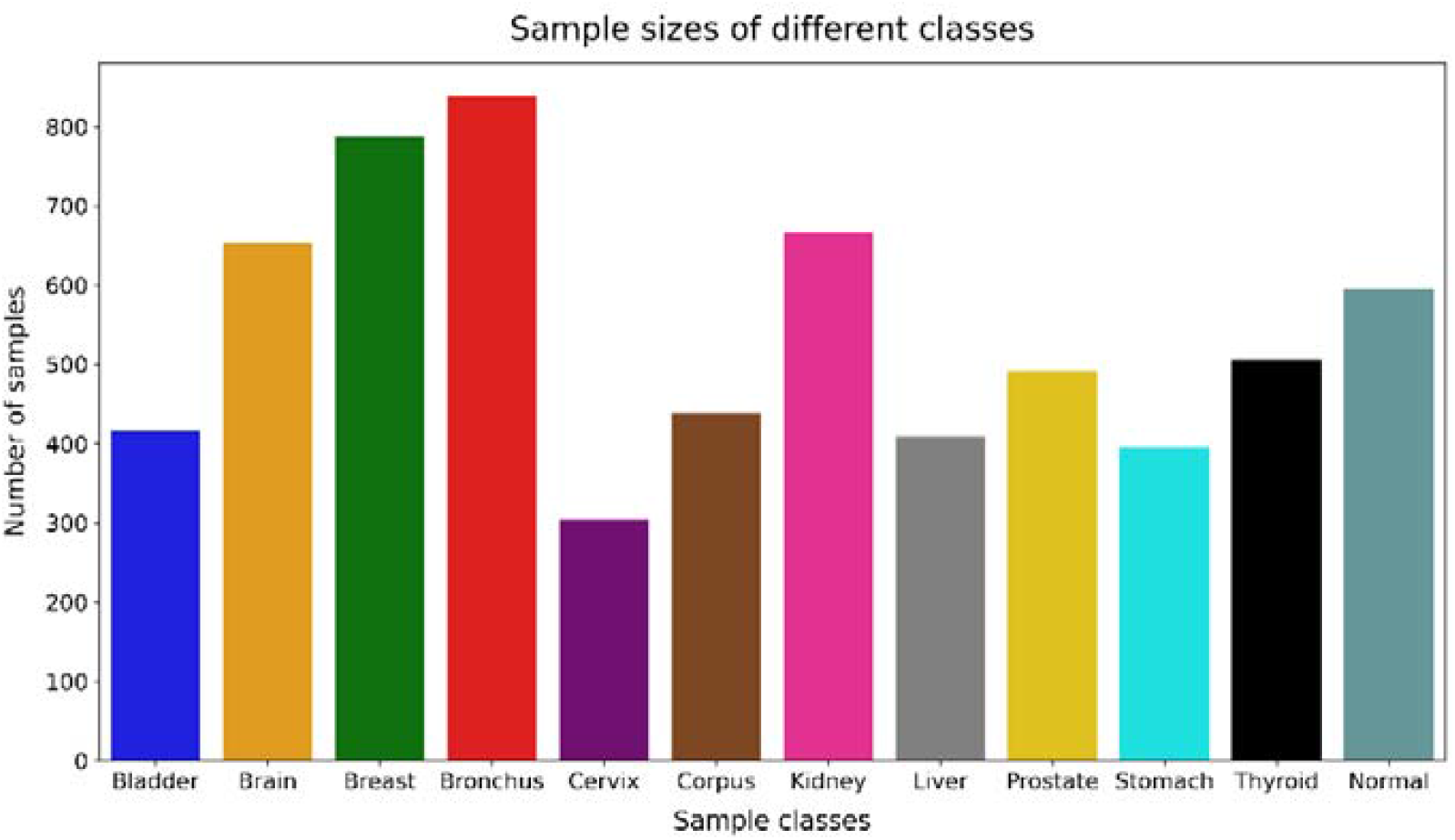
The distribution of TCGA samples.

**supplementary Figure S2.**
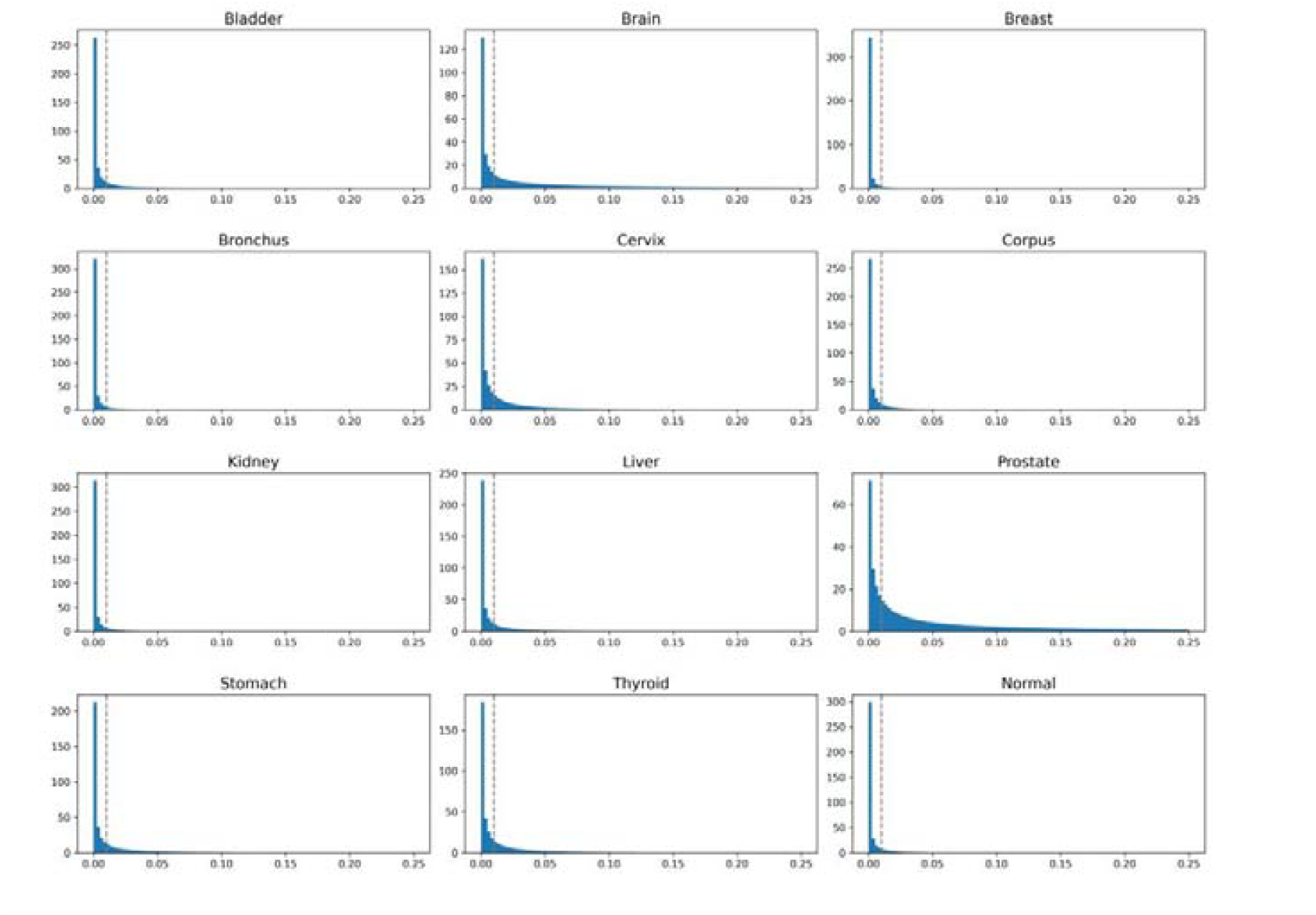
The weight distribution in each network.

**supplementary Figure S3.**
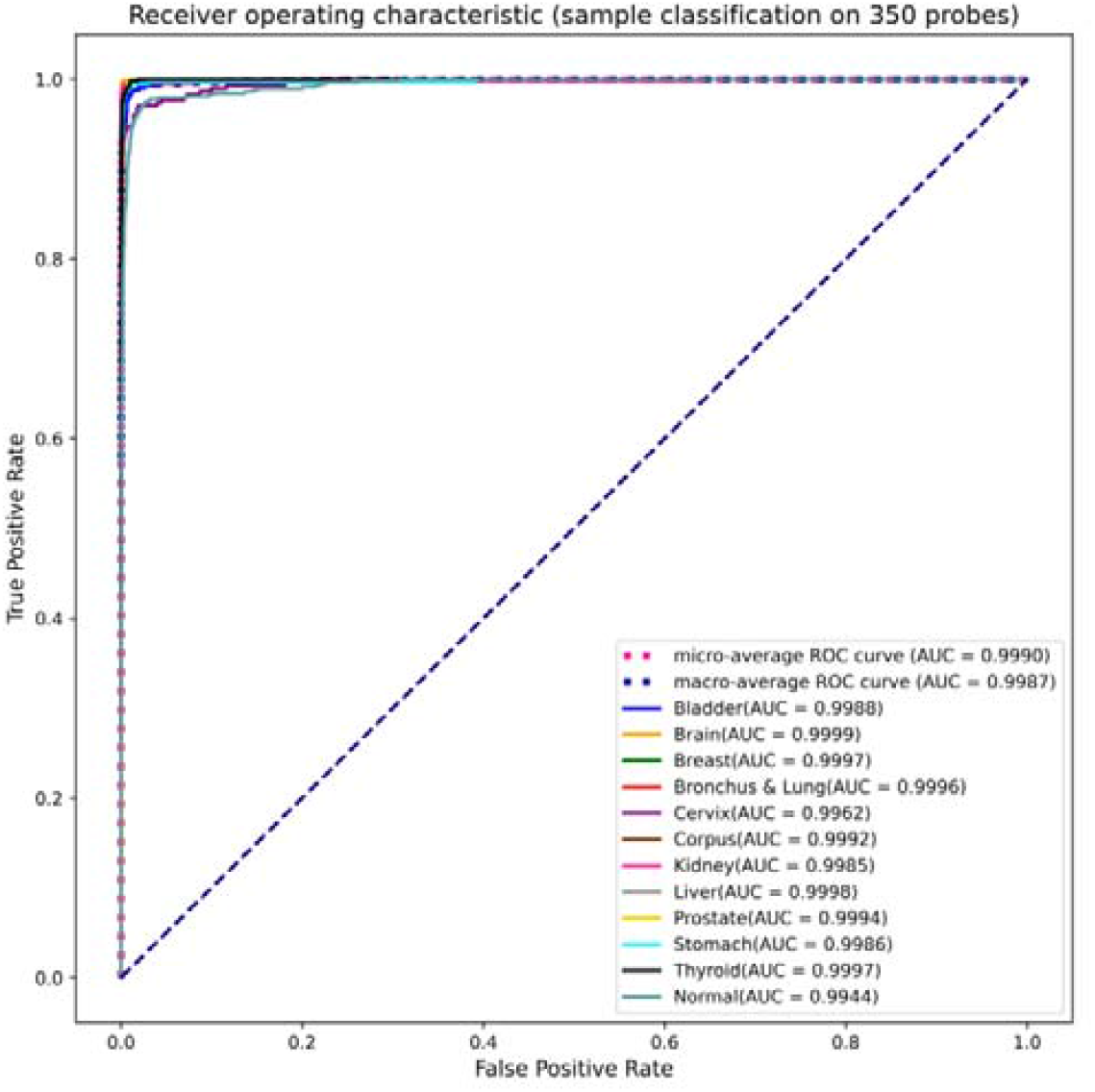
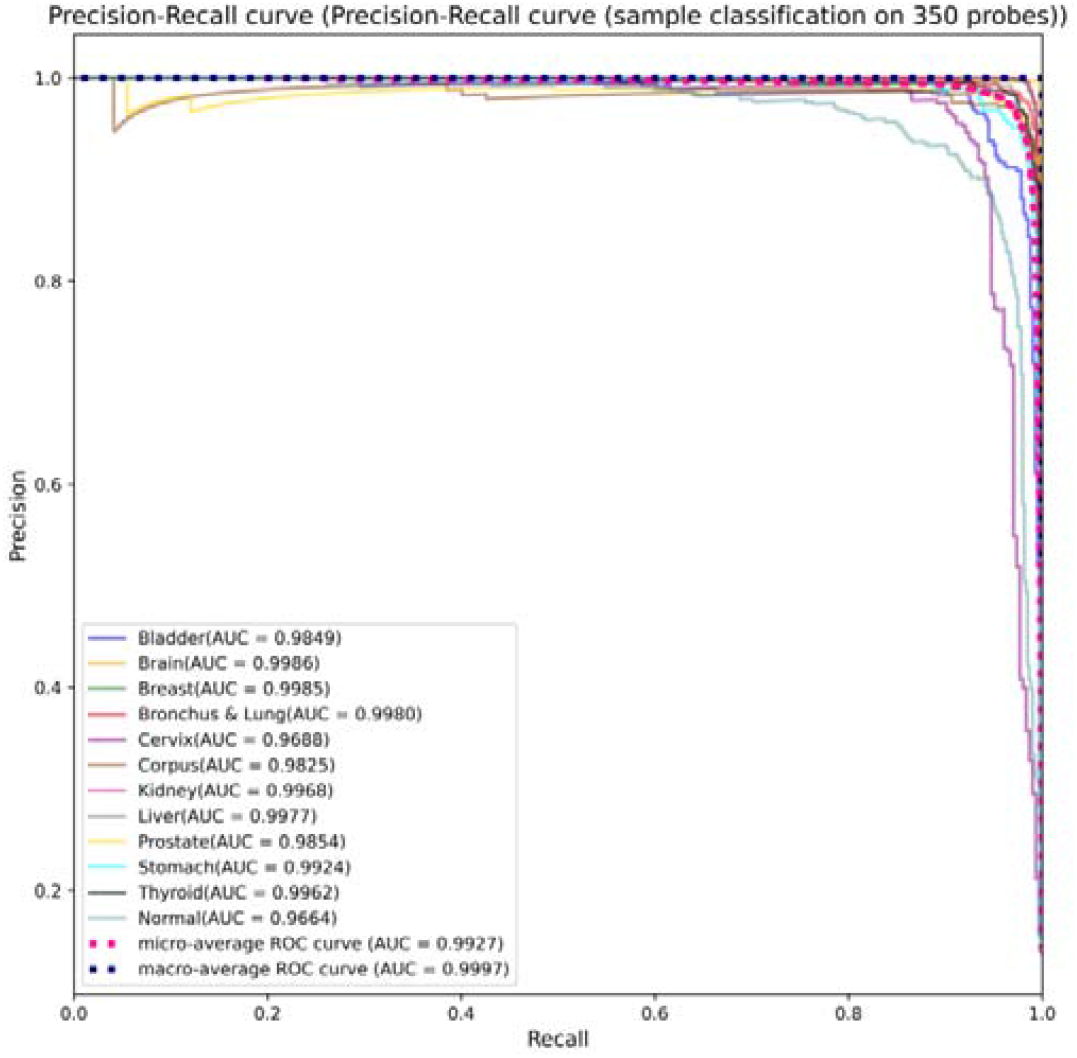
The ROC and PR curves of SAGCN model with *k*_3_ = 0.3 on the TCGA dataset.

**supplementary Figure S4.**
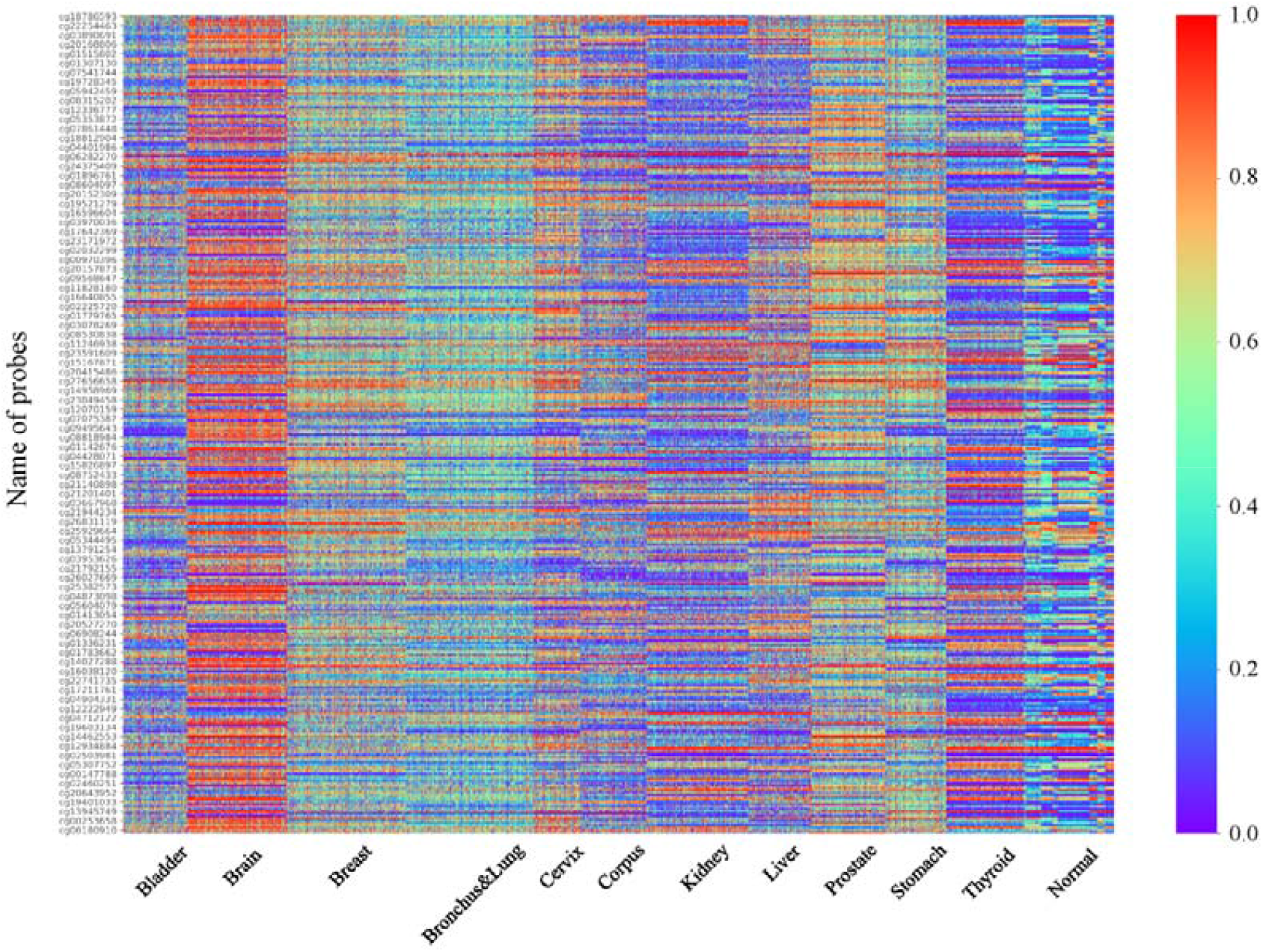
The cluster heatmap of the selected DEMs.

