## Supplementary material for "A Self-attention Graph Convolutional Network for Precision Multi-tumour Early Diagnostics with DNA Methylation Data": Table

Table 1. The performance of SAGCN with different
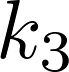
 value

| 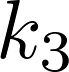 | fold | Macro-average | | | | | | | |
| --- | --- | --- | --- | --- | --- | --- | --- | --- | --- |
|  |  | ME | BS | LL | Recall | Precision | F1-score | AUC | AUPR |
| 0.1 | avg | 0.0533 | 0.0065 | 0.0230 | 0.9467 | 0.9538 | 0.9441 | 0.9963 | 0.9953 |
|  | std | 0.0640 | 0.0065 | 0.0212 | 0.0640 | 0.0546 | 0.0672 | 0.0044 | 0.0047 |
| 0.2 | avg | 0.0417 | 0.0051 | 0.0184 | 0.9583 | 0.9621 | 0.9571 | 0.9983 | 0.9975 |
|  | std | 0.0236 | 0.0029 | 0.0106 | 0.0236 | 0.0222 | 0.0242 | 0.0018 | 0.0019 |
| 0.3 | avg | 0.0383 | 0.0060 | 0.0213 | 0.9617 | 0.9631 | 0.9613 | 0.9971 | 0.9965 |
|  | std | 0.0510 | 0.0076 | 0.0263 | 0.0510 | 0.0500 | 0.0515 | 0.0049 | 0.0049 |
| 0.4 | avg | 0.0233 | 0.0031 | 0.0128 | 0.9767 | 0.9793 | 0.9766 | 0.9990 | 0.9980 |
|  | std | 0.0232 | 0.0034 | 0.0139 | 0.0232 | 0.0198 | 0.0233 | 0.0012 | 0.0022 |
| 0.5 | avg | **0.0083** | **0.0013** | **0.0049** | **0.9917** | 0.9918 | **0.9917** | 0.9998 | 0.9995 |
|  | std | 0.0105 | 0.0014 | 0.0044 | 0.0105 | 0.0103 | 0.0105 | 0.0002 | 0.0006 |
| 0.6 | avg | 0.0083 | 0.0014 | 0.0053 | 0.9917 | **0.9924** | 0.9916 | **0.9999** | **0.9997** |
|  | std | 0.0053 | 0.0006 | 0.0020 | 0.0053 | 0.0048 | 0.0053 | 0.0001 | 0.0003 |
| 0.7 | avg | 0.0133 | 0.0026 | 0.0125 | 0.9867 | 0.9884 | 0.9867 | 0.9995 | 0.9986 |
|  | std | 0.0135 | 0.0026 | 0.0109 | 0.0135 | 0.0117 | 0.0136 | 0.0005 | 0.0012 |

Table 2. The sample classification performance of SAGCN + SVM with different
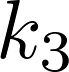
 value

| 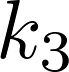 | fold | Macro-average | | | | | | | |
| --- | --- | --- | --- | --- | --- | --- | --- | --- | --- |
|  |  | ME | BS | LL | Recall | Precision | F1-score | AUC | AUPR |
| 0.1 | avg | 0.0871 | 0.0115 | 0.0441 | 0.9110 | 0.9111 | 0.9099 | 0.9929 | 0.9985 |
|  | std | 0.0043 | 0.0005 | 0.0016 | 0.0045 | 0.0039 | 0.0038 | 0.0005 | 0.0001 |
| 0.2 | avg | 0.0511 | 0.0068 | 0.0273 | 0.9466 | 0.9454 | 0.9455 | 0.9969 | 0.9989 |
|  | std | 0.0041 | 0.0003 | 0.0008 | 0.0037 | 0.0037 | 0.0034 | 0.0002 | 0.0001 |
| 0.3 | avg | 0.0417 | 0.0055 | 0.0223 | 0.9559 | 0.9553 | 0.9551 | 0.9979 | 0.9991 |
|  | std | 0.0030 | 0.0003 | 0.0011 | 0.0026 | 0.0036 | 0.0031 | 0.0002 | 0.0001 |
| 0.4 | avg | 0.0391 | 0.0051 | 0.0209 | 0.9582 | 0.9578 | 0.9577 | 0.9981 | 0.9991 |
|  | std | 0.0030 | 0.0003 | 0.0008 | 0.0036 | 0.0033 | 0.0034 | 0.0001 | 0.0001 |
| 0.5 | avg | 0.0374 | 0.0049 | 0.0200 | 0.9593 | 0.9599 | 0.9592 | 0.9983 | 0.9992 |
|  | std | 0.0023 | 0.0003 | 0.0008 | 0.0012 | 0.0036 | 0.0024 | 0.0001 | 0.0001 |
| 0.6 | avg | 0.0343 | 0.0045 | 0.0186 | 0.9631 | 0.9628 | 0.9626 | 0.9985 | 0.9993 |
|  | std | 0.0037 | 0.0003 | 0.0009 | 0.0035 | 0.0046 | 0.0041 | 0.0001 | 0.0001 |
| 0.7 | avg | **0.0301** | **0.0040** | **0.0170** | **0.9668** | **0.9675** | **0.9669** | **0.9987** | **0.9993** |
|  | std | 0.0034 | 0.0003 | 0.0009 | 0.0037 | 0.0046 | 0.0042 | 0.0001 | 0.0001 |

Table 3. The classification performance of SAGCN + SVM for each kind tumors and normal samples, respectively

| Sample | fold | Macro-average | | | | | | | |
| --- | --- | --- | --- | --- | --- | --- | --- | --- | --- |
|  |  | ME | BS | LL | Recall | Precision | F1-score | AUC | AUPR |
| Bladder | avg | 0.0062 | 0.0051 | 0.0205 | 0.9456 | 0.9590 | 0.9520 | 0.9989 | 0.9857 |
|  | std | 0.0019 | 0.0012 | 0.0037 | 0.0203 | 0.0218 | 0.0134 | 0.0006 | 0.0061 |
| Brain | avg | 0.0008 | 0.0009 | 0.0054 | 0.9950 | 0.9967 | 0.9958 | 0.9999 | 0.9987 |
|  | std | 0.0008 | 0.0008 | 0.0032 | 0.0067 | 0.0040 | 0.0046 | 0.0001 | 0.0014 |
| Breast | avg | 0.0026 | 0.0025 | 0.0122 | 0.9862 | 0.9924 | 0.9892 | 0.9998 | 0.9987 |
|  | std | 0.0010 | 0.0010 | 0.0034 | 0.0084 | 0.0046 | 0.0042 | 0.0002 | 0.0013 |
| Bronchus | avg | 0.0058 | 0.0045 | 0.0194 | 0.9761 | 0.9785 | 0.9773 | 0.9996 | 0.9979 |
|  | std | 0.0009 | 0.0005 | 0.0015 | 0.0040 | 0.0047 | 0.0037 | 0.0003 | 0.0010 |
| Cervix | avg | 0.0062 | 0.0050 | 0.0219 | 0.9361 | 0.9327 | 0.9339 | 0.9965 | 0.9711 |
|  | std | 0.0013 | 0.0008 | 0.0028 | 0.0155 | 0.0322 | 0.0123 | 0.0017 | 0.0061 |
| Corpus | avg | 0.0037 | 0.0035 | 0.0152 | 0.9754 | 0.9705 | 0.9729 | 0.9992 | 0.9837 |
|  | std | 0.0011 | 0.0012 | 0.0042 | 0.0102 | 0.0048 | 0.0074 | 0.0005 | 0.0142 |
| Kidney | avg | 0.0032 | 0.0020 | 0.0113 | 0.9836 | 0.9847 | 0.9841 | 0.9986 | 0.9970 |
|  | std | 0.0015 | 0.0007 | 0.0025 | 0.0114 | 0.0056 | 0.0073 | 0.0017 | 0.0026 |
| Liver | avg | 0.0031 | 0.0026 | 0.0109 | 0.9537 | 0.9973 | 0.9749 | 0.9998 | 0.9976 |
|  | std | 0.0014 | 0.0009 | 0.0024 | 0.0184 | 0.0055 | 0.0111 | 0.0001 | 0.0015 |
| Prostate | avg | 0.0046 | 0.0036 | 0.0141 | 0.9691 | 0.9693 | 0.9691 | 0.9994 | 0.9873 |
|  | std | 0.0018 | 0.0015 | 0.0053 | 0.0198 | 0.0131 | 0.0126 | 0.0005 | 0.0112 |
| Stomach | avg | 0.0046 | 0.0035 | 0.0149 | 0.9701 | 0.9566 | 0.9629 | 0.9989 | 0.9934 |
|  | std | 0.0016 | 0.0012 | 0.0044 | 0.0242 | 0.0185 | 0.0111 | 0.0017 | 0.0056 |
| Thyroid | avg | 0.0031 | 0.0031 | 0.0126 | 0.9797 | 0.9803 | 0.9800 | 0.9997 | 0.9961 |
|  | std | 0.0014 | 0.0011 | 0.0034 | 0.0097 | 0.0111 | 0.0095 | 0.0002 | 0.0028 |
| Normal | avg | 0.0165 | 0.0119 | 0.0459 | 0.9309 | 0.8925 | 0.9110 | 0.9946 | 0.9669 |
|  | std | 0.0018 | 0.0009 | 0.0039 | 0.0156 | 0.0251 | 0.0139 | 0.0018 | 0.0079 |

Table 4. The classification performance of SAGCN +SVM for tumors of stage I

| Sample | Macro-average | | | | | | | | | | | | | | |
| --- | --- | --- | --- | --- | --- | --- | --- | --- | --- | --- | --- | --- | --- | --- | --- |
|  | ME | BS | | LL | | Recall | | Precision | | F1-score | | | AUC | | AUPR |
| Bladder | 0.0022 | | 0.0022 | | 0.0097 | | 1.0000 | | 0.6667 | | 0.8000 | 0.9992 | | 0.8354 | |
| Breast | 0.0000 | | 0.0002 | | 0.0030 | | 1.0000 | | 1.0000 | | 1.0000 | 1.0000 | | 1.0000 | |
| Bronchus | 0.0100 | | 0.0081 | | 0.0272 | | 0.9534 | | 1.0000 | | 0.9761 | 1.0000 | | 0.9999 | |
| Kidney | 0.0056 | | 0.0070 | | 0.0282 | | 0.9823 | | 1.0000 | | 0.9911 | 1.0000 | | 1.0000 | |
| Liver | 0.0000 | | 0.0009 | | 0.0044 | | 1.0000 | | 1.0000 | | 1.0000 | 1.0000 | | 1.0000 | |
| Stomach | 0.0078 | | 0.0057 | | 0.0274 | | 0.9799 | | 1.0000 | | 0.9899 | 0.9991 | | 0.9989 | |
| Thyroid | 0.0011 | | 0.0007 | | 0.0033 | | 1.0000 | | 0.6667 | | 0.8000 | 1.0000 | | 1.0000 | |

Table 5. The classification performance of SAGCN +SVM for tumors of stage II

| Sample | Macro-average | | | | | | | | | | | | | | |
| --- | --- | --- | --- | --- | --- | --- | --- | --- | --- | --- | --- | --- | --- | --- | --- |
|  | ME | BS | | LL | | Recall | | Precision | | F1-score | | | AUC | | AUPR |
| Bladder | 0.0154 | | 0.0140 | | 0.0451 | | 0.9542 | | 1.0000 | | 0.9766 | 0.9999 | | 0.9998 | |
| Breast | 0.0103 | | 0.0046 | | 0.0167 | | 1.0000 | | 0.5000 | | 0.6667 | 1.0000 | | 1.0000 | |
| Bronchus | 0.0077 | | 0.0075 | | 0.0257 | | 0.9684 | | 1.0000 | | 0.9840 | 0.9999 | | 0.9998 | |
| Kidney | 0.0051 | | 0.0028 | | 0.0131 | | 0.9615 | | 1.0000 | | 0.9804 | 1.0000 | | 1.0000 | |
| Liver | 0.0000 | | 0.0002 | | 0.0029 | | 1.0000 | | 1.0000 | | 1.0000 | 1.0000 | | 1.0000 | |
| Stomach | 0.0026 | | 0.0016 | | 0.0105 | | 1.0000 | | 0.9865 | | 0.9932 | 1.0000 | | 1.0000 | |
| Thyroid | 0.0051 | | 0.0029 | | 0.0127 | | 0.9655 | | 0.9655 | | 0.9655 | 0.9998 | | 0.9977 | |

Table 6. The performance comparison of SAGCN + SVM with classic machine learning methods

| Method |  | Macro-average | | | | | | | |
| --- | --- | --- | --- | --- | --- | --- | --- | --- | --- |
|  | fold | ME | BS | LL | Recall | Precision | F1-score | AUC | AUPR |
| DT | avg | 0.1154 | 0.0178 | 0.5749 | 0.8777 | 0.8777 | 0.8769 | 0.9388 | 0.6713 |
|  | std | 0.0041 | 0.0006 | 0.0310 | 0.0041 | 0.0056 | 0.0038 | 0.0024 | 0.0082 |
| RF | avg | **0.0275** | 0.0086 | 0.0420 | **0.9687** | **0.9710** | 0.9696 | 0.9982 | 0.9991 |
|  | std | 0.0029 | 0.0001 | 0.0004 | 0.0021 | 0.0039 | 0.0031 | 0.0004 | 0.0001 |
| ERT | avg | 0.0297 | 0.0082 | 0.0402 | 0.9658 | 0.9703 | 0.9677 | 0.9982 | 0.9991 |
|  | std | 0.0037 | 0.0001 | 0.0004 | 0.0033 | 0.0039 | 0.0035 | 0.0003 | 0.0001 |
| GNB | avg | 0.1054 | 0.0175 | 0.5837 | 0.8930 | 0.9012 | 0.8954 | 0.9775 | 0.6901 |
|  | std | 0.0083 | 0.0014 | 0.0453 | 0.0108 | 0.0071 | 0.0088 | 0.0029 | 0.0194 |
| SAGCN  + SVM | avg | 0.0301 | **0.0040** | **0.0170** | 0.9668 | 0.9675 | **0.9669** | **0.9987** | **0.9993** |
|  | std | 0.0034 | 0.0003 | 0.0009 | 0.0037 | 0.0046 | 0.0042 | 0.0001 | 0.0001 |

Table 7. The performance comparison of SAGCN + SVM with the state-of-the-art methods

| Method |  | Macro-average | | | | | | | |
| --- | --- | --- | --- | --- | --- | --- | --- | --- | --- |
|  | fold | ME | BS | LL | Recall | Precision | F1-score | AUC | AUPR |
| CGATCPred | avg | 0.0309 | 0.0044 | 0.0287 | 0.9660 | 0.9676 | 0.9666 | 0.9982 | 0.9718 |
|  | std | 0.0042 | 0.0005 | 0.0031 | 0.0051 | 0.0046 | 0.0049 | 0.0004 | 0.0177 |
| DeepCDR | avg | 0.0237 | 0.0032 | 0.0161 | 0.9749 | 0.9747 | 0.9746 | 0.9989 | 0.9990 |
|  | std | 0.0024 | 0.0004 | 0.0025 | 0.0032 | 0.0028 | 0.0030 | 0.0003 | 0.0002 |
| SAGCN  +SVM | avg | 0.0301 | 0.0040 | 0.0170 | 0.9668 | 0.9675 | 0.9669 | 0.9987 | **0.9993** |
|  | std | 0.0034 | 0.0003 | 0.0009 | 0.0037 | 0.0046 | 0.0042 | 0.0001 | 0.0001 |

Table 8. The performance evaluation of SAGCN + SVM with external validation dataset of tissue DNA methylation

| Dataset | Tumour | Macro-average | | | | | | | |
| --- | --- | --- | --- | --- | --- | --- | --- | --- | --- |
|  | type | ME | BS | LL | Recall | Precision | F1-score | AUC | AUPR |
| GSE90496 | Brain | 0.1464 | 0.1189 | 0.3567 | 0.7789 | 1.0000 | 0.8757 | 1.0000 | 1.0000 |
| GSE155207 | Kidney | 0.0023 | 0.0052 | 0.0269 | 0.9500 | 1.0000 | 0.9744 | 1.0000 | 1.0000 |
| GSE158075 | Bronchus & Lung | 0.0586 | 0.0371 | 0.1284 | 0.8378 | 0.6078 | 0.7045 | 0.9689 | 0.8028 |
| GSE164988 | Stomach | 0.0248 | 0.0151 | 0.0567 | 1.0000 | 0.4762 | 0.6452 | 0.9910 | 0.5376 |
| TCGA | Normal | 0.2050 | 0.1153 | 0.3817 | 0.5902 | 0.3529 | 0.4417 | 0.8271 | 0.5563 |

Table 9. The performance evaluation of SAGCN + SVM with cell free DNA methylation data

| Dataset | Tumour | Macro-average | | | | | | | |
| --- | --- | --- | --- | --- | --- | --- | --- | --- | --- |
|  | type | ME | BS | LL | Recall | Precision | F1-score | AUC | AUPR |
| GSE157272 | Prostate | 0.0068 | 0.0042 | 0.0144 | 0.8636 | 1.0000 | 0.9268 | 1.0000 | 1.0000 |

supplementary Table 1. The detailed patient demographics and clinical characteristics of the TCGA data set

supplementary Table 2. The extracted DEMs by SAGCN and the related annotations

supplementary Table 3 The classification performance of SAGCN for the 11 type tumors, respectively

| Sample | fold | Macro-average | | | | | | | |
| --- | --- | --- | --- | --- | --- | --- | --- | --- | --- |
|  |  | ME | BS | LL | Recall | Precision | F1-score | AUC | AUPR |
| Bladder | avg | 0.0033 | 0.0038 | 0.0139 | 0.9800 | 0.9800 | 0.9800 | 0.9998 | 0.9981 |
|  | std | 0.0067 | 0.0038 | 0.0133 | 0.0400 | 0.0400 | 0.0400 | 0.0004 | 0.0038 |
| Brain | avg | 0.0000 | 0.0000 | 0.0001 | 1.0000 | 1.0000 | 1.0000 | 1.0000 | 1.0000 |
|  | std | 0.0000 | 0.0000 | 0.0002 | 0.0000 | 0.0000 | 0.0000 | 0.0000 | 0.0000 |
| Breast | avg | 0.0000 | 0.0005 | 0.0021 | 1.0000 | 1.0000 | 1.0000 | 1.0000 | 1.0000 |
|  | std | 0.0000 | 0.0008 | 0.0026 | 0.0000 | 0.0000 | 0.0000 | 0.0000 | 0.0000 |
| Bronchus | avg | 0.0033 | 0.0030 | 0.0112 | 0.9800 | 0.9800 | 0.9800 | 0.9996 | 0.9965 |
|  | std | 0.0067 | 0.0034 | 0.0101 | 0.0400 | 0.0400 | 0.0400 | 0.0007 | 0.0070 |
| Cervix | avg | 0.0000 | 0.0000 | 0.0001 | 1.0000 | 1.0000 | 1.0000 | 1.0000 | 1.0000 |
|  | std | 0.0000 | 0.0000 | 0.0002 | 0.0000 | 0.0000 | 0.0000 | 0.0000 | 0.0000 |
| Corpus | avg | 0.0000 | 0.0000 | 0.0006 | 1.0000 | 1.0000 | 1.0000 | 1.0000 | 1.0000 |
|  | std | 0.0000 | 0.0000 | 0.0005 | 0.0000 | 0.0000 | 0.0000 | 0.0000 | 0.0000 |
| Kidney | avg | 0.0033 | 0.0021 | 0.0076 | 0.9800 | 0.9800 | 0.9800 | 0.9996 | 0.9960 |
|  | std | 0.0067 | 0.0037 | 0.0104 | 0.0400 | 0.0400 | 0.0400 | 0.0007 | 0.0080 |
| Liver | avg | 0.0050 | 0.0045 | 0.0155 | 0.9800 | 0.9618 | 0.9705 | 0.9991 | 0.9890 |
|  | std | 0.0076 | 0.0057 | 0.0187 | 0.0400 | 0.0468 | 0.0398 | 0.0014 | 0.0176 |
| Prostate | avg | 0.0000 | 0.0000 | 0.0000 | 1.0000 | 1.0000 | 1.0000 | 1.0000 | 1.0000 |
|  | std | 0.0000 | 0.0000 | 0.0000 | 0.0000 | 0.0000 | 0.0000 | 0.0000 | 0.0000 |
| Stomach | avg | 0.0017 | 0.0012 | 0.0036 | 0.9800 | 1.0000 | 0.9895 | 1.0000 | 1.0000 |
|  | std | 0.0033 | 0.0024 | 0.0065 | 0.0400 | 0.0000 | 0.0210 | 0.0000 | 0.0000 |
| Thyroid | avg | 0.0000 | 0.0000 | 0.0001 | 1.0000 | 1.0000 | 1.0000 | 1.0000 | 1.0000 |
|  | std | 0.0000 | 0.0000 | 0.0001 | 0.0000 | 0.0000 | 0.0000 | 0.0000 | 0.0000 |
| Normal | avg | 0.0000 | 0.0005 | 0.0038 | 1.0000 | 1.0000 | 1.0000 | 1.0000 | 1.0000 |
|  | std | 0.0000 | 0.0004 | 0.0024 | 0.0000 | 0.0000 | 0.0000 | 0.0000 | 0.0000 |
